# Identification of post-transcriptional modifications in nucleic acid sequences using purpose-designed molecular beacons

**DOI:** 10.1101/2023.05.25.542360

**Authors:** Parul Sahu, Getulio Pereira, Jian Wang, Kamil Szeliski, Nikolay Dokholyan, Vivian Tran, Miroslaw Korneck, John Tigges, Jennifer Jones, Ionita Ghiran

## Abstract

Post-transcriptional RNA modifications (PTxMs) present in small RNA species, specifically circulating extracellular RNAs, were recently identified as clinically relevant readouts, often more indicative of disease severity than the classical “up and down” changes in their copy number alone. While identification of PTxMs requires multiple and complex sample preparation steps, microgram-range amounts of RNA, followed by expensive and protracted bioinformatics analyses, the clinically relevant information is usually a yes/no for a particular genetic variant(s), and an up/down answer for relevant biomarkers. We have previously shown that molecular beacons (MBs) can identify specific nucleic acid sequences with picomolar sensitivity and single nucleotide specificity by exploiting the target-dependent change in their electrophoretic mobility profile. We now present a method for direct identification of miRNAs and isomiRs in cells and extracellular vesicles using gel electrophoresis, without the need for RNA isolation and purification. The detection is based on discreet changes in the hydrodynamic surface profile, the overall size, charge and charge distribution of the MB-target hybrid. Furthermore, using an RNA tertiary structure prediction algorithm (iFoldRNA) and a custom molecular dynamics simulation (DMD), we designed modified MBs specific for m6A-modified nucleotides in target RNA sequences. The sample preparation method coupled to the software package affords the design of specific MBs and sensitive, multiplex-type detection of targets in a wide variety of biofluids and cells, in a simple mix and read approach.

## Introduction

Molecular beacons (MBs) are hairpin-shaped oligonucleotides (RNA or DNA) that contain a dedicated anti-sense hybridization sequence to a specific single-stranded RNA or DNA molecule, a double-stranded, short stem region, and at their termini, a fluorochrome, and a quencher situated within Foster’s radius for an effective quencing.^1,2,3,4^ In the absence of the target, the stem sequence brings the quencher into proximity to the fluorochrome, preventing it from fluorescing upon excitation. The binding of the MB to the complementary target by its hybridization sequence triggers a conformational change in the beacon, opening the stem and separating the quencher from the fluorochrome, allowing emission of fluorescence following excitation. We have previously shown that gel electrophoresis offers an alternate MB-based readout method for identification of specific oligonucleotide sequences and differentiate between RNA and DNA molecules with identical sequences, solely based on their electrophoretic pattern. The method, which reaches picomolar sensitivity and single nucleotide specificity was tested using synthetic oligonucleotides as targets and validated on total RNA isolated from blood cells, such as RBCs and platelets.

In the last decade, circulating cell-free nucleic acids^5-i7^ present either in extracellular vesicles (EVs)^8-10^, or associated RNA- and DNA-binding proteins have received increased attention not just for their critical role in basic biological processes, but also as potential theragnostic agents for a growing number of cancers.^11-13^ One of the challenges facing precision oncology is the discovery and the validation of reliable biomarkers, followed by the development of sensitive, multiplex sets of detection tests. Overcoming these limitations would allow quantification of patterns of clinical significance within the biomarker panels with profound, direct, and life-saving outcomes for patient care. Quantitative detection of various species of cell-free nucleic acids is usually performed using laborious and time-consuming methods, such as qPCR, next-generation sequencing, microarray-based hybridization, or northern blotting. More recently, MBs,^2,4^ have started to be used successfully in clinical settings for high sensitivity miRNA-based molecular diagnosis of cancer.^3, 14^ Efficient hybridization of molecular probes to the intended sequence found in biofluids requires a mix of three distinctive, yet converging purpose buffers, which dissociate the nucleic acids from their RNA/DNA-binding proteins without altering their structure, reduce the free energy required for MB-target hybridization, while maintaining the specificity of the reaction, and promote molecular crowding, forcing the MBs and targets closer together increasing the sensitivity of detection. These new class of sample buffers offer the promise of rapid target detection, bypassing altogether the need for time consuming nucleic acid extraction steps^(1-2)^.

Most DNA/RNA-based responses to stress or disease are centered, in addition to the canonical quantitative, up/down-type changes in their expression levels, to reversible, qualitative modification of specific nucleosides. These modifications are promoted by a set of specialized proteins with opposing roles, the Writers, which add specific chemical groups to certain nucleotides in specific sequences, and the Erasers, which removed the modifications. The majority of the nucleotide modifications involve either the attachment of a chemical group, usually methyl or hydroxy-methyl at a specific position either on the base such as, adenine, (m^1^A) or (m^6^A), cytosine, (m^3^C) or (m^5^C), guanosine (m^1^G), (m^7^G), the ribose sugar (e.g., 2′-*O*-methyladenosine), or in some instances on both base and sugar such as *N*^6^,2′-*O*-dimethyladenosine (m^6^Am). The overall altered sequence is then recognized by dedicated RNA/DNA binding proteins named Readers, whose function is then changed based on the type and location of the modification on the molecule.^15-18^ Functionally, posttranscriptional modifications (PTxMs) of RNA molecules represent a second layer of information which conveys subtle, transient, location-dependent cues modulating the RNA message. More recently, PTxMs found in circulating exRNAs, are seen as promising markers in an ever-increasing number of cancers. Changes in cytosine methylation (m^5^C), of several mature circulating miRNAs species were reported to be positively correlated with poor prognosis in certain cancers, whereas others with increased survival rate.^(3)^

Identification of these post-transcriptional RNA modifications is primarily performed by one of four separate approaches: immunoprecipitation, mass spectrometry, direct sequencing, and reverse transcriptase, respectively ^(4-5)^. All these methods, require large amounts of isolated nucleic acids, usually in the microgram range, and use involved bioinformatics analyses and approaches to identify specific base modifications either as bulk or with limited base resolution.

A second type of post-transcriptional modification underwent by miRNAs is represented by the addition, subtraction or editing of bases controlled by exo-ribonucleases, nucleotidyl transferase and ADAR (adenosine deaminase acting on dsRNA) enzymes, respectively. Initially considered sequencing errors and dismissed as such, in the last decade their existence and role in cancers evolution were started to be understood, when experimental indicating that isomiRs derived from the same hairpin can target distinct sets of mRNA, due to changes in the seed sequence ^(6)^.

We here describe a new method for identification of cellular and extracellular vesicle RNA molecules containing PTxMs, using purposefully designed molecular beacons and gel electrophoresis. This approach uses a dedicated lysis and hybridization buffer along with modified MBs designed specifically for detection of m6A nucleotides in target sequences, and isoforms on miR451 present in RBCs and RBC-EVs. The basis of the approach relies on discreet changes in the hydrodynamic surface profile, and the overall charge and charge location of the MB-target hybrid in the presence of epigenetic modifications. These conformational changes of the MB-target complex directly and measurably influence its electrophoretic mobility compared to the identical, unmodified sequence. The new MB-target conformation promoted by the presence of methylation groups on nucleic acids and molecular beacons were simulated using machine learning-based algorithms and validated using synthetic oligos, and RNA from cells and EVs collected from human donors. We show here using a simple mix and read approach, that isomiRs and hydroxymethyl cytosine-modified sequences are detectable by wild type MB, and in the case of m6A, by modified MBs.

## Material and Methods

### Reagents

Dulbecco’s phosphate-buffered saline (DPBS(1X), without calcium, or magnesium, 2.6 mM KCl, 1.47 mM KH2 PO4, 137 mM NaCl, and 8.05 mM Na2HPO4), was obtained from Thermo Fisher Scientific (Waltham, MA). Novex™ TBE Running Buffer (5X), and Novex TBE Gels, 20% were obtained from Thermo Fisher Scientific. Gel Loading Dye, purple (5X), without SDS was obtained from New England Biolabs (Ipswich, Massachusetts). Hybridization buffer was a gift from Abcam (Waltham, MA).

### Molecular beacons and targets sequences

Molecular beacons, synthetic miRNAs or DNA/RNA oligonucleotide analogs were obtained from Integrated DNA technologies IDT (Coralville, IA) or Genelink (Elmsford, NY). MBs were conjugated with a 5’ end either fluorochrome 1, (FL1) or fluorochrome 2 (FL2). The PTxM miRNAs or MBs had hydroxy-C or A, or m6A modification at the basis noted in the figures. All the MBs and corresponding target sequences used for this project are shown in **Table 1**.

**Table 1.**
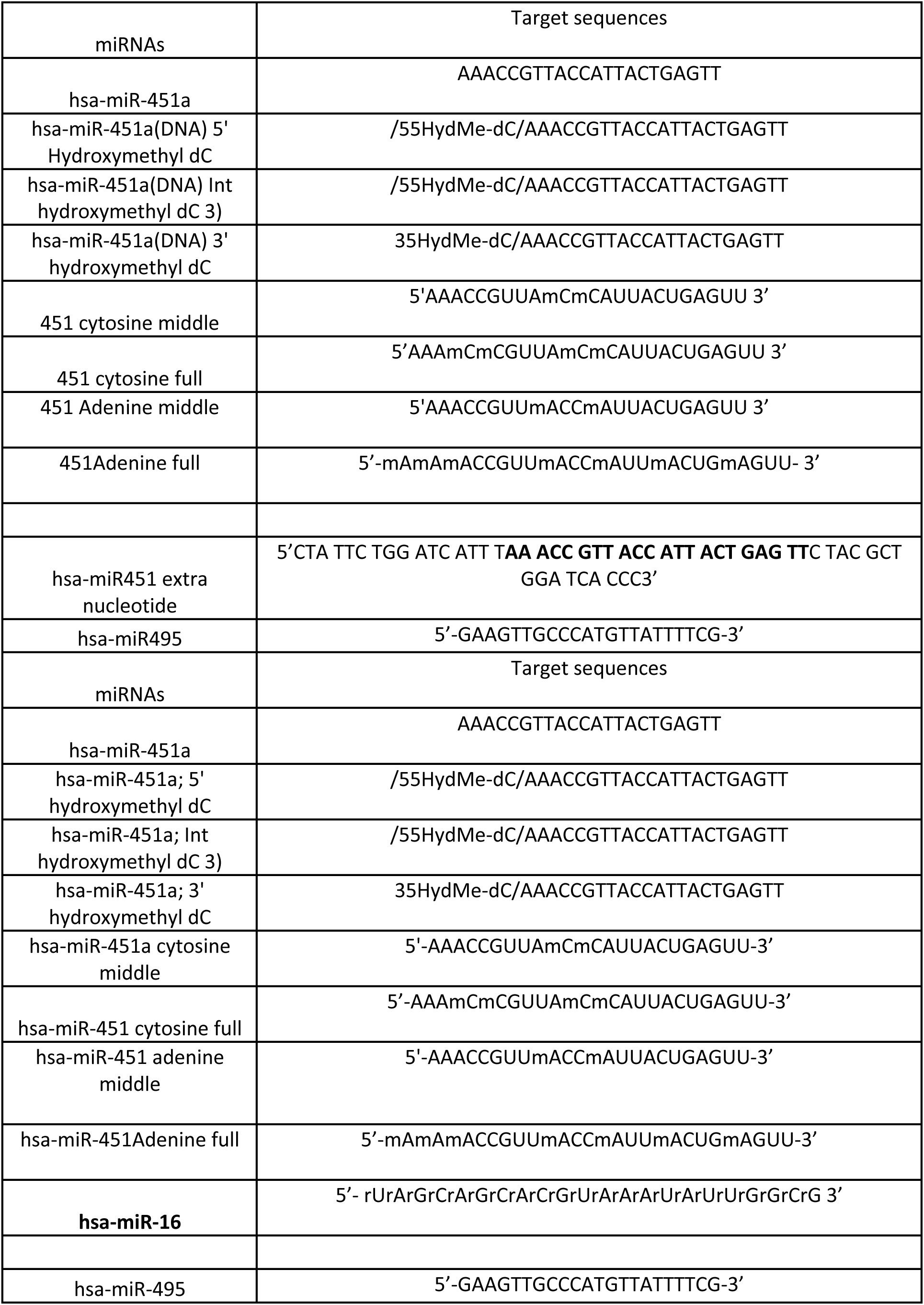
Target RNAs sequences.

### Fluorometric measurement of miRNA-MB hybridization

MBs were diluted in different concentrations (10% and 25%) of the hybridization buffer or 100 μL of dPBS1X to a final concentration of 50 nM, and then incubated with 50 nM concentrations of synthetic miRNA target analog in 96 well plates (Corning™ 96-Well clear bottom, black walls) for 40 min at 37 °C. The fluorescence intensity of each well was measured (λEx 495 nm; λEm 521 nm) by a microplate reader (Synergy HT Multi-Mode, Biotek, Winooski, VT, USA). For the kinetics assay, MBs were diluted in 100 μL of 1X dPBS or 10% and 25% of the hybridization buffer to a final concentration of 50 nM, and then incubated with either 50 nM target analog or 50 nM mismatch sequences in 96 well plates. Fluorescence readings were acquired at 37°C every 5 min using a Promega fluorometer.

### miRNA-MB hybridization detection by gel electrophoresis

After incubation of the molecular beacons with or without miRNAs target analogs for 40 min at 37°C, 12uL of each sample was mixed with Gel Loading dye (2.5uL), and then loaded on Novex TBE 20% gels. Gel electrophoresis was performed with constant voltage for 10 min at 70 V, and then the voltage was increased to 130 V for an additional 40 min. The MB fluorescence signal was visualized a ChemiDoc MP Imaging System (Bio-Rad, Hercules, CA). Exposure times were set on “Manual” and varied depending on the sample between 10 and 120s.

### RNA isolation from cell free plasma

Blood samples were collected from two self declared healthy donors and plasma was isolated by sequentially three step centrifugation at 500xg, 2500xg and 12000xg for 10 mins. RNA was isolated by using miRNeasy Serum/Plasma Kit (Qiagen, Germantown, MD). The concentration of isolated RNA was measured by using Qubit™ microRNA Assay Kit in a Qubit 4 Fluorometer (Thermo Fisher, Waltham, MA).

### Generation of extracellular vesicles

The collection of blood was performed under a study which was approved by the Beth Israel Deaconess Medical Center Institutional Review Board (IRB #2001P000591). Blood was obtained via venipuncture using Vacutainer EDTA tubes (BD, Franklin Lakes, NJ) from self-declared healthy volunteers. Platelet-rich plasma was removed by centrifugation at 500 x g for 10 min. The pellet was diluted in 1x PBS, and contaminating leukocytes were removed by passing the blood through an Acrodisc WBC syringe filter (Pall Corporation, NY). RBCs were washed by centrifugation at 500xg for 5 mins twice in HBSS^++^. One hundred microliters of washed RBCs resuspended in 5 mL in HBSS^++^ were then incubated with 10 micromolar of ionomycin on a slow shaker at RT at 37C for 60 minutes. The cells were spun down at 10,000xg and the supernatant containing RBC-EVs concentrated using 10k cut off Amicon tubes. The resulting EVs were characterized by dark field and nano-flow cytometry.

### Machine learning

The MD simulations were performed using the *pmemd.cuda* program within the Amber 18 software package. Previous studies have shown that certain versions of Amber force fields yield satisfactory results when investigating the stability and dynamics of RNAs. For this study, we utilized the Amber RNA OL3 force field. The starting structures were generated using iFoldRNA and then prepared via the teLeap module in Amber 18. The nucleotides were methylated using PyMol, and the parameters for these modifications were obtained from the Amber RNA OL3 force field. To solvate the RNA molecules, we employed an octahedral box containing TIP3P water molecules, with the RNA atoms maintaining a distance of 9Å from the box boundary.

Given that metal ions play a crucial role in the stability and dynamics of RNA, we added potassium ions to neutralize the system, followed by additional K+ and Cl-ions. The concentration of potassium ions and chloride ion in our simulation box is approximately 0.3 M. To ensure the reliability of our MD simulations, we conducted several energy minimization steps prior to the actual simulation. Specifically, we first performed energy minimization of the entire system using the steepest descent method for 3000 steps, followed by an additional 3000 steps using the conjugate gradient method. Next, we used harmonic positional restraints to maintain the positions of the RNA atoms and solely minimized the positions of water and ions. Once this was complete, we performed energy minimization of the entire system once again. During the MD simulation phase, we initially applied weak restraints to the RNA molecules and allowed the system to heat up from 0 K to our target temperature of 300 K over the course of 200 ps. Once this heating process was complete, we removed the restraints on the RNA molecules and carried out explicit solvent MD under constant pressure. The length of hydrogen bonds was constrained using the shake algorithm, and we employed the isothermal-isobaric (NPT) ensemble to maintain a constant temperature of 300 K and pressure of 1 bar throughout the simulation. The time step was set to 2 fs.

## RESULTS

### Hybridization buffer increases the sensitivity of the target detection in a sequence and concentration dependent manner

Hybridization buffers promote crowding of molecules in solution by limiting the amount of solvent between the molecules increasing the chances of encounter, allowing the interaction between molecules to occur with higher efficacy, effectively increasing the local concentration of macromolecules (**Fig. 1A**), mimicking the transient microdomains present observed in living cells^(7)^. We tested the effect of the hybridization buffer on the efficiency of the MBs binding with its targets, by first measuring the fluorescence and the kinetics of the reaction using fluorometry. Fifty nM MBs (**Fig. 1B**) were incubated with 50 nM miRNAs target analogs or mismatched controls in the presence of standard loading buffer (black diamond), or 10% (triangle) or 25% (star) hybridization buffer added to the loading buffer. The fluorescence signal was then recorded every 5 min for a total experimental time of 2 hrs. The fluorescence peak was achieved between 30 and 40 min depending on the RNA target, with higher concentration of hybridization buffer in the sample rendering higher fluorescence signals. Based on the fluorometry results, we next tested whether the hybridization buffer would improve the fluorescence signal when using gel electrophoresis. The incubation time for the MB-target was chosen to match the maximum intensity signal obtained in fluorometry and based on our previous work^(8)^. We tested the effect of the hybridization buffer by using a concentration of the target which was at or below the detection limit when using standard loading buffer. We incubated 1pM of miR-451a MB and miR16-5p-MB with same concentrations of their miRNAs target analogs, or mismatched controls, in buffer alone, or increasing concentrations of hybridization buffer (1%, 10% and 25%). The intensity of the fluorescence bands representative of the MB-target hybridization, increased with the concentration of the hybridization buffer (**Fig. 1C**), paralleling the results obtained using fluorometry, albeit at a significantly lower concentration. Increasing the concentration of the hybridization buffer at 50% did not increased the intensity of the signal above that obtained at 25% (**Fig. 1D**), suggesting that at a given density of analytes, the efficiency of the buffer diminishes. Testing higher concentrations of the hybridization buffer by gel electrophoresis was not possible, due to significant distortion in the migration pattern (data not shown).

**Fig. 1.**
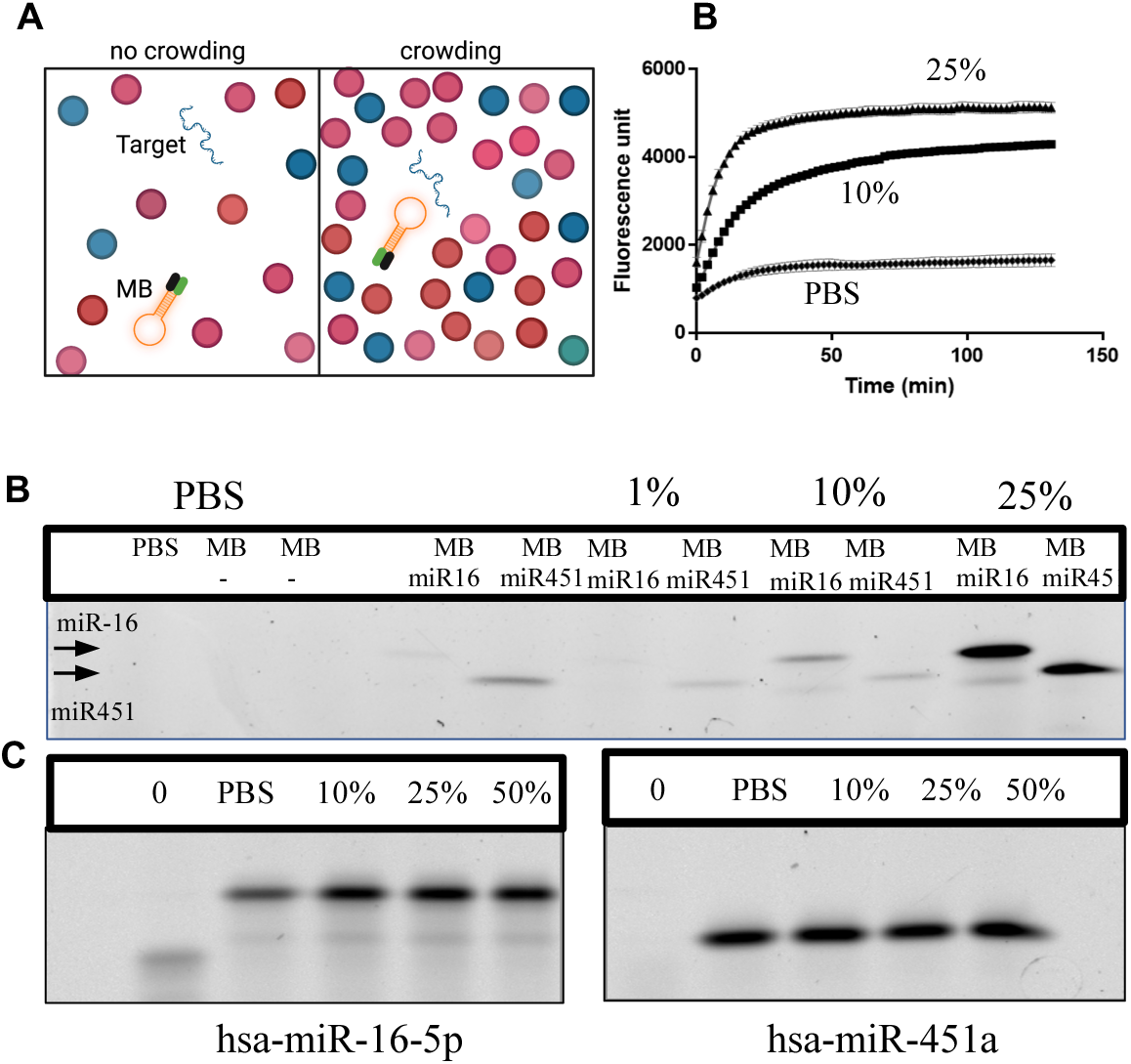
Hybridization buffer effect diminishes with increase target concentration. **A**) Schematics of the effect molecular crowding agents have on the kinetics of the reaction. **B**. Fifty nM miR541-a were incubated with the target or in the presence of standard buffer (diamond), 10 % (square) or 25% (triangle) hybridization buffer and analyzed by fluorometry. **C**) One pM miR451-a and 16-5p were incubated in the presence of standard buffer or increasing concentrations of hybridization buffer. **D)** The experiment was repeated using 50 nM targets. Samples were incubated for 15 mins and separated on 20% TBE gel.

### Post-transcriptional modifications alter the affinity and the fluorescence intensity generated by MBs

Recently, epigenetic modifications of either the genomic DNA or circulating extracellular RNAs, such as methylations, in particular m6A, as well as hydroxy-methylations, of the nitrogenous base or the corresponding sugar backbone, were described in certain cancers, and showed to provide enhance the functional context for disease progression than the classical up/down changes in their concentrations alone(^3, 9^). Previously, we have shown that the location of the mismatch nucleotide between the MB and the target is important for the formation, fluorescence, and electrophoretic characteristics of the resulting hybrid. Therefore, we next generated targets with modified nucleotides, in addition to the 5’ end, also in the middle of the sequence and at 3’ end (**Fig. 2**). The results show that the electrophoretic delay was dependent on the location of the modified nucleotide, with the modification located at the 3” promoting (see **Table 1**) the largest electrophoretic delay. When we performed the competition experiment, where the wt- and the modified target were mixed at the same concentration, the separation between the two hybrids was maintained, suggesting that the approach can be used with samples where there is an expectation of encountering both states. (**Fig. 2A last lane**). Densitometric data of the bands obtained in lane 6 indicate that the binding kinetics of the MB to the hydroxymethyl oligos is favored compared to the wild type, thus rendering a stronger band, although when probes individually, lanes 4 and 5 respectively, the intensity of the bands were virtually identical. The electrophoretic mobility of the MB-hydroxy-methylated target was slower than that of the MB-wt target, suggesting that small changes in the surface area and the shape of the MB-target hybrid may have a significant effect on its electrophoretic behavior.

**Fig. 2.**
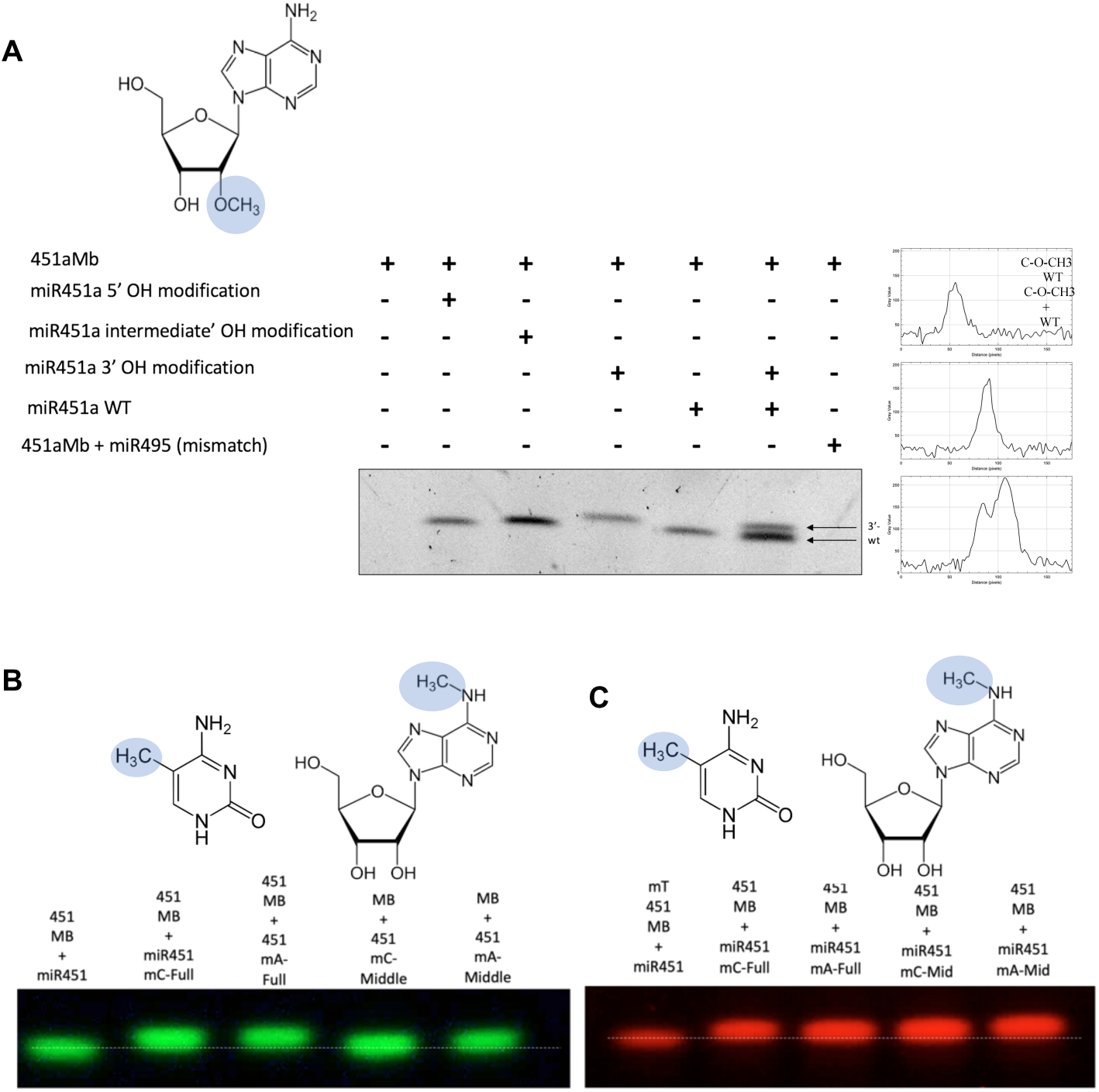
Standard MBs identify the 2’-O-methyl-modified RNA sequences. **A)**. MB-miR-451a were incubated with wild type and hydroxyl-methylated targets prior to gel electrophoresis and the fluorescence recorded by a fluorescence imager (Geldoc, Biorad). The modifications at the 3” end rendered a better separation than those found at 5’ end or in the middle of the miRNA (arrows). **B** and **C)**. Standard MBs do not reliably identified methylated A or C residues. Wt, m6A, and m5C (Table 1) were probed either with st-MB or ms-MBs, modified at the corresponding U or G respectively, but failed to render distinct pattern.

### The standard molecular beacon design fails to identify m6A nucleotides

The most widespread modifications found in nucleic acids, during both normal and pathological conditions are the N6-methyl adenosine (m6A) and 5-methyl cytosine (m5C) with the former especially common in tRNA, tRNA fragments, miRNAs and other small RNA species ^(10-11)^. We repeated the experiment delineated in **Fig. 2A**, using as targets miR451-a with A replaced with m6A at the middle A (see Table 1, line 5). Our data indicate that although the modified oligo induced an electrophoretic delay compared to the wild type oligos, the differences were minimal (**Fig. 2B**), even when the concentration of the gel was increased from 20% to 30% (data not shown). As a first step to overcome this problem, we methylated the corresponding U residue on the MBs to generate a MB-target hybrid with a significantly altered surface profile, in the hope of promoting a measurable change in its electrophoretic mobility. The results in **Fig. 2C**, showed that the new m3U-MB difference in the migration pattern of the wt vs m6A oligo was still minimal.

### Discrete molecular dynamics simulations help understand the effect of fluorochromes/quencher pairs and nucleobase modifications on the electrophoretic mobility of MBs

To design and test MB with electrophoretic ability to identify and differentiate between m6A modified sequences, we began by performing molecular dynamics simulations to computationally visualize the effect of several fluorochromes/quencher pairs and altered nucleobases on the conformation of the MB alone and MBs following hybridization with various targets. The MBs which were generated using modified nucleobases and specific fluorochromes/quenchers were named *<u>m</u>*ethylation *<u>s</u>*ensitive-MB, or ms-MB, whereas the *<u>st</u>*andard MBs, or st-MB. The overall workflow employed is shown in **Fig. 3A** and detailed in the ***Material and Methods*** section. The results of the simulation indicated that in addition to modifying the nucleobase corresponding or near the methylated nucleotide, an additional change had to be added to the structure of the beacon to generate a beacon sensitive enough to differentiate between a methylated and non-methylated nucleobases. Fluorochromes are a diverse group photoreactive compounds with a wide range of sizes and charges, which once attached to a protein or nucleic acid can significantly alter the conformation and behavior of their substrates ^(12)^. The goal of the simulation was to identify a MB which could differentiate between sequence identical wt and m6A nucleic acids, not just by a slow/fast electrophoretic signature, but rather through another parameter which would be independent by the small variation of the expected length of miRNAs, such as isomiRs. The molecular simulation results (**Fig. 3B**) indicated that in the case of wt-MB there was a stable conformation, which represented the majority (70.6%) of the possible forms, whereas for ms-MB, the simulation identified 3 clusters of stable conformations (0.293 and 0.288 for clusters 1 and 2, and 0.205 for cluster 3 respectively). Following simulations, we compared the electrophoretic pattern of the ms-MB to the wt-MBs, by initially running them on a gel without targets, to test whether the stable conformations predicted by the simulation were matching the electrophoretic pattern. While in the MB close conformation the quenching of the fluorochromes by the quencher approaches 98%^(13)^, at high concentrations of 200 nM and above, and exposure times over 5 minutes, the location of the closed MB in the gel is detectable, as we have previously shown^(8)^. The resulting electrophoretic patterns of closed MBs show that while the wt-MB formed only one band (**Fig. 3C**, left), the candidate for ms-MB formed two distinct bands and a smear (**Fig. 3C**, right), suggesting the simulation results were extrapolatable, at least in part, to the electrophoretic behaviors of the MBs. We next contrasted the gel electrophoretic profile with the data obtained by simulated conformations and energy landscape generated by the interaction of the two MBs with wt or m6A modified miRNA-451a.

**Fig. 3.**
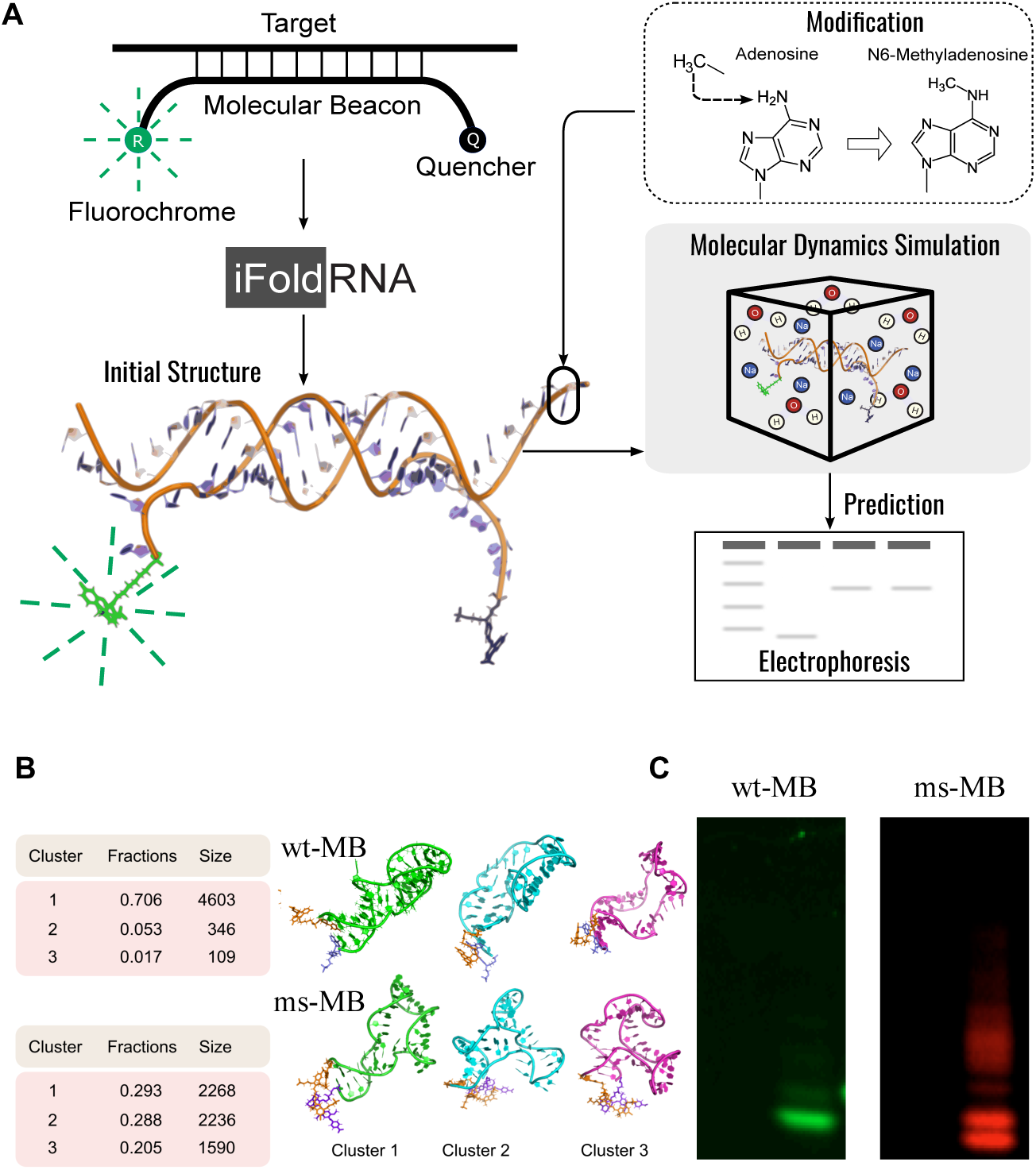
Discrete molecular dynamics simulations illustrate the differences between molecular beacons using different dyes. **A)**. The computational workflow used to test the MB and MB-target fold. An initial three-dimensional structure of the MB/target complex with was generated by iFoldRNA. Modifications were introduced using PyMol, and molecular dynamics simulations tests were run for the MB alone, target alone and the MB/target complex, followed by calculations the thermodynamics properties to project their performance in electrophoresis. **B)**. Simulation predict the electrophoretic behavior. The three largest clusters of the frames in the simulation trajectory for miR451-wt (a) and miR451-ms (b). **C)**. The type of fluorochrome and modified nucleobase alter the molecular conformation of MBs. Wt-(left gel) and ms-MB (right gel) were run in the absence of targets at a concentration of 200nM and exposed for 6 mins to record the location of folded MBs.

### The electrophoretic mobility of the MB-targets depends on the methylation status of RNA, as well as the composition of the MB

We began by generating 2-D and 3-D conformations of the MB-target complexes to visualize the effect of the added methyl groups (highlighted in blue) on adenine and the corresponding uracil, respectively. The gray line labeled with an asterisk, indicates the shift in the angle of the bond between the ribose and nucleobase of the modified pair, compared to the un-modified nucleobases. The effects of the new conformation on the 3-D structure in shown in **Fig. 4B**. These results suggested that the alteration if the pairing of the based due to the presence of methyl groups on both, the target as well as the beacon, may generate a complex which may be able to differentiate between methylated and unmethylated target. We next performed four groups of simulations of the energy levels of the hybrids, as follows: st-MB/Target, st-MB/m6A-target, ms-MB/target, and ms-MB/m6A-target. We completed 500 ns MD simulations, and then plotted the free energy landscape for each combination. As shown in **Fig. 4C** and **D**, in the case of st-MB interacting with either wt or methylated oligos, the free energy distribution was nearly centered, suggesting that the MB would run as a uniform complex, and be less likely to differentiate between the two targets. In the case of ms-MB (**Fig. 4E** and **F**), the interaction with wt or m6A modified targets, respectively, resulted in a dispersed free energy landscape that could be grouped into several distinct areas, indicating that the structure may adopt one of the several stable conformations. The results of the simulation also indicated that the complexes may not move freely from one conformation to another, due to the high energy barriers between different conformations, showed as light red bridges, labeled with asterisk. Furthermore, for ms-MBs incubated with either the wildtype target or the m6A target, there is one cluster (Cluster 1 in **Fig. 4E** and **F**) that has a similar sizes (RG ≈ 20.8 Å, vs. 20.71 Å) in both cases, suggesting that one of the possible conformations of the ms-MB may behave similarly when hybridizing with either wt or m6A target. However, Clusters 2 had a different RG sizes (22.05Å vs. 21.7 Å) upon binding the wt or m6A target, predicting that these clusters may display a different electrophoretic behavior. In agreement with our simulations, when incubated with wt- or m6A-targets, st-MBs rendered one band **(Fig. 4 G**, lanes 1 and 2), whereas the complex of ms-MB with either wild type (lane 3), m6A (lane 4) or both forms (lane 6), resulted in two main bands, and a minor one. The intensity and location of the top, strong bands (**Fig. 4G**, labeled “m6A insensitive”) were indicative of size of the target (electrophoretic speed) and amount (fluorescence intensity), but not methylation status, as the top bands in lanes 3 (wt) and 4 (m6A) were at the same level. In turn, the faster band was sensitive to amount (intensity) and methylation status of the target (location), being able to differentiate between *wt* and m6A miR451 either alone (**Fig. 4G**, lane 3) and (**Fig. 4G**, lane 4) or mixed (**Fig 4G**, lane 6). **Fig 4H** shows the histograms corresponding to the white vertical lines, across methylation sensitive bands on lanes 3 (a), 4(b) and 6 (c), which represent the locations where the line profiles (side histograms) were calculated. The lower panel in **Fig. 4G** focuses on the methylation sensitive bands obtained in a separate experiment where the red channel was exposed for 70 seconds instead of 15 (top). Thus, a combination of modified MB and a specific fluorochrome with a unique charge/size is necessary to generate a methylation sensitive MB able to detect and quantify: **i**) the amount of target (fluorescence intensity), **ii**) its size (electrophoretic drag), and **iii**) its methylation level expressed as a ratio of the AUC of the wt and m6A line profiles. We have also investigated whether the methylation sensitive MB will bind to the wt and M6A targets with the same affinity, by performing a competition experiment. Wild type and methylated oligos were incubated with the methylation sensitive MBs at ratios ranging from 1:1 to 1:10. The results in **Fig 4I** show that the methylation sensitive beacon had a similar affinity for both forms, being able to discriminate between samples with various concentrations of wt and m6A modified targets.

**Fig. 4.**
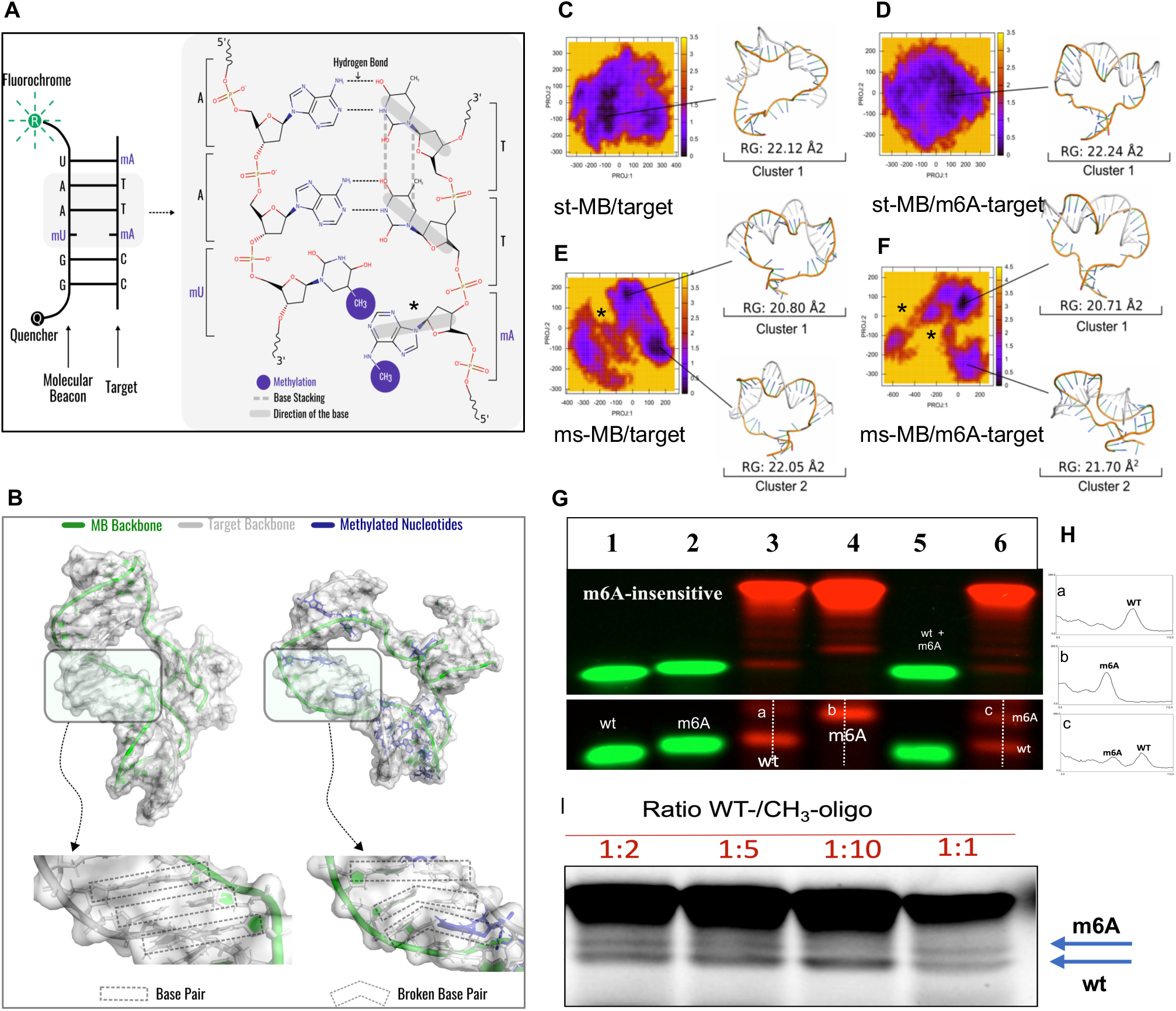
Identification on m6A modifications depends on both, beacon methylation as well as the choice of fluorochrome. **A)**. Schematic representation of the effect methylation of A and corresponding U has on the 2-D structure of the MB-target hybrid. **B)**. 3-D modeling of the MB-target hybrid focusing on the methylated nucleobases. Gel electrophoresis of methylated beacon at -MB with wild type (lane 3), methylated (lane 4) or mix targets (lane 6) coupled to wt-fluorochrome or ms-fluorochrome. The methylation insensitive band is labeled m6A-insensitive. **C**-**F)**. The free energy landscapes of wt-MB/Target, ms-MB/Target, wt-/mTarget, and ms-MB-/mTarget. Target is the miRNA-451, and mTarget is the m6A miRNA-451. See results for details. **G)**. Electrophoretic patterns recapitulate simulation results. Wt-MB and ms-MBs were incubates with wt-(lanes 1 and 3) or m6A-(lanes 2 and 4) or mixed together (lanes 5 and 6). In a separate experiment, the lower bands were exposed longer and densitometry analysis performed as described in Methods. **H)**. Intensity profiles of the methylation specific bands were calculated along the lines labels “a”, “b” and “c” and shown in histogram panel on the right). **I**. Modified MB bind wt and m^6^A targets equally effective. Ms-MB was incubated with a mix of wt or m6A modified miR451 at ratios shown and resolved using 20% gel.

### 1.5. Direct detection of RNAs in cells and extracellular vesicles by gel electrophoresis

An efficient lysis buffer aimed at isolating RNA from a sample of biological material, should release most of the existing RNA in sample, free of any contaminants likely to inhibit or interfere with processing or downstream analyses. We tested efficiency of the lysis buffer in detection EV-RNA using RBC-EVs generated following our published protocols ^(14-15)^. The presence of RBC-EVs in cell supernatant was confirmed by high resolution, double immersion dark field microscopy (**Fig. 5A**, left, DF) and negative staining TEM (**Fig. 5A**, right, EM), The EVs further characterized by in Nanoflow cytometry (**Fig. 5B**), the concentration determined by Exoid (Izon, Cambridge MA, data not shown). RBC-EVs, at a concentration of 10^7^/lane were then incubated with either st- or ms-MB in the presence of lysis buffer. Although miR-451 contains a canonical DRACH motive (AA^m^**A**CC) in its sequence^(16)^, our results show that the methylation sensitive MB did not identify a subpopulation of methylated miR451 in RBC-EVs, which, if successful should have generated a double band below the main methylation insensitive band, similar to that in **Fig 5C**. These results suggest that miR451 in RBC-EVs is either not m6A methylated despite the presence of the DRACH motif, or the sensitivity of our approach was insufficient to generate any detectable signal provided by a miR451 methylated subpopulation.

**Fig. 5.**
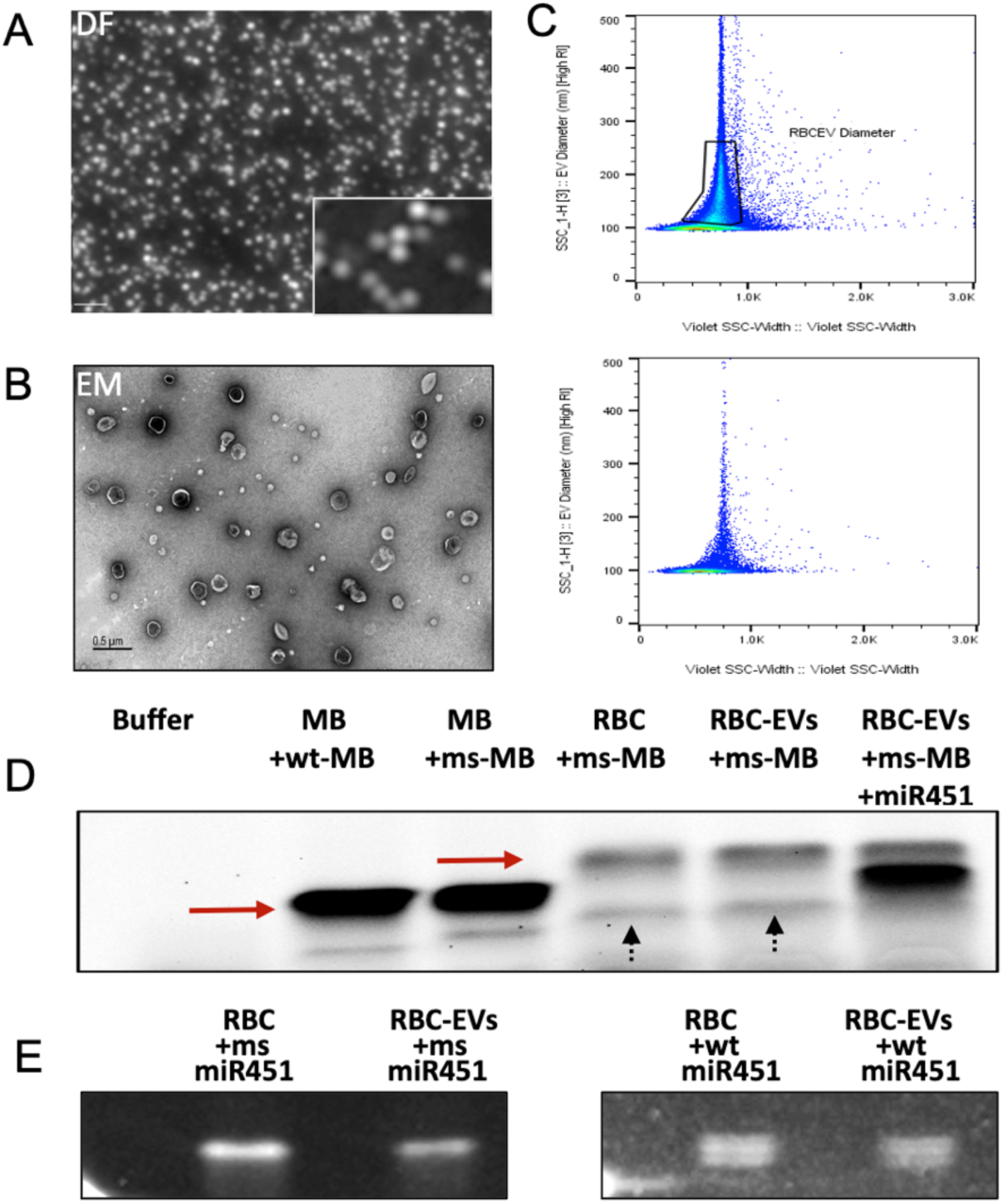
Direct identification of specific miRNA from RBCs and RBC-EVs. Characterization of RBC-derived EVs. Double immersion dark field microscopy (**Fig. 6A**, DF,) images with a 40×0.95 UplaXApo objective. Inset detailed of the field images using an 100×1.35 iris UplanApo objective. Bar 1um. TEM negative staining (**Fig. 6B**, right, EM) of isolated EVs showing two population of EVs. **C**) RBC-EVs were analyzed by nanoflow cytometry The results are shown as dotplots and are displayed in nm on the *Y*-axis, and violet scatter (SSC) values on the *X*-axis. The gate (RBC-EV Diameter) represents the area were particles below 300 nm are located. **D**) Direct identification of miR451 in RBCs and RBC-EVs. RBCs and RBC-EVs were incubated with lysis and hybridization buffer for 10 mins prior to adding the ms-MBs. Lane 1 represents MB alone, lanes 2, positive controls, MB451 spiked with miRNA 451 *wt* and miR451*m6A* oligos respectively. Lanes 4-5, 10^5^ RBCs isolated from ah healthy donor and 10^7^ RBC-EVs incubated with lysis buffer and ms-MB, showing expected miR451 bands. Arrows point to bands generated by the miR-451a MB fluorescent hybrid indicating the lack of detectable m6A-modified miR-451a. **E**). RBC and RBC-EVs contain isomiR451. RBC and RBC-EVs were incubated with ms-MBs (left) and wt-MBs (right) in the presence of hybridization and lysis buffer. Unlike ms-sensitive MB, wt-MBs identify isoforms of miR451 with one nucleotide difference (see Table 3).

### MB electrophoresis identifies discreet changes in miRNA sequence length

MicroRNA isoforms (isomiRs) are a heterogenous family of variable length, nucleotide composition, or both compared to the parent miRNA. The range of isomiRs length varies in between -4 to +8 nt compared to the canonical form. While most miRNAs are expressed almost ubiquitously, certain isomiR species are cell- or tissue-depended^(17)^. The tissue and gender specificity and differential expression is more pronounced for isomiRs than the parent miRNAs^(18)^. We next tested the ability of the MBs to detect the presence of isomiRs in RBCs and RBC-derived EVs by direct lysis of the cells and EV with sample buffer and incubation with MBs. Our results in **Fig. 5E** show what both RBCs and RBC EVs contain at several forms of miR451 with different sequence lengths thus rendering distinct miR-451 positive bands. Somehow surprising, the ms-MBs did not have the spatial resolution to identify the presence of isoforms with few different nucleotide, unlike the st-MBs. Our RNA-seq analysis identified multiple isomiR451-a with only few having the ability to bind the MB synthesized against the parent miRNA-451a. In Table 3, sequences labeled with an asterisk are likely the ones identified by the st-MB, having only one nt difference at the 3’ end from the canonical sequence. The seed sequence of the isomiRs is underscored. The electrophoretic mobility of the isomiRs were similar in both the RBCs and EVs suggesting a lack of sorting mechanism, at least based on the size separation afforded by our MB (**Fig 5E)**. Our RNA-seq data shown that RBCs (Table 1) contain several isomiR-451 with extra nucleotides at either at the 5’ or 3’ end as previously reported, and lengths varying between 24 and 17 nt, compared to the parent miR-451a with 22 nt^(19)^. The lack of identification of additional isomiRs of miR451-a is likely due to high specificity of the MB, which reaches single nucleotide resolution, but only if the mismatched base is located inside the sequence and not at one of the termini, as others and we have shown have previously shown(^8, 20-21)^.

**Table 2.**
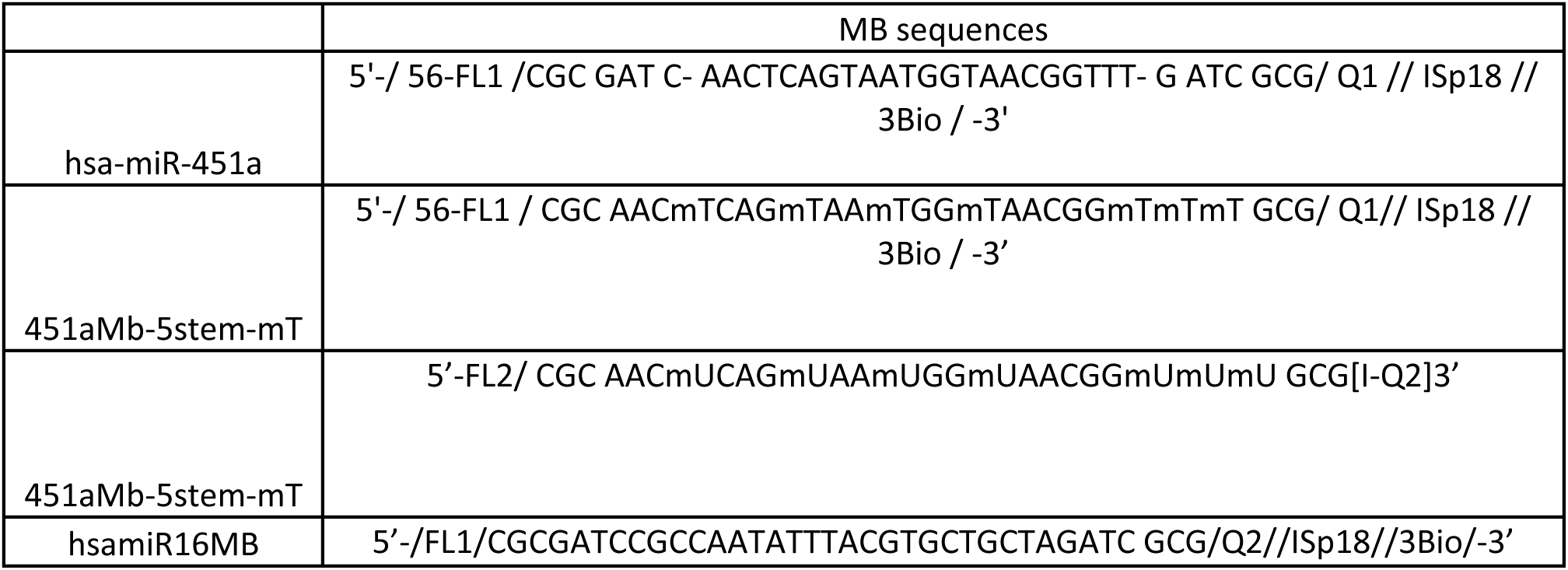
Molecular beacon sequences.

**Table 3.**
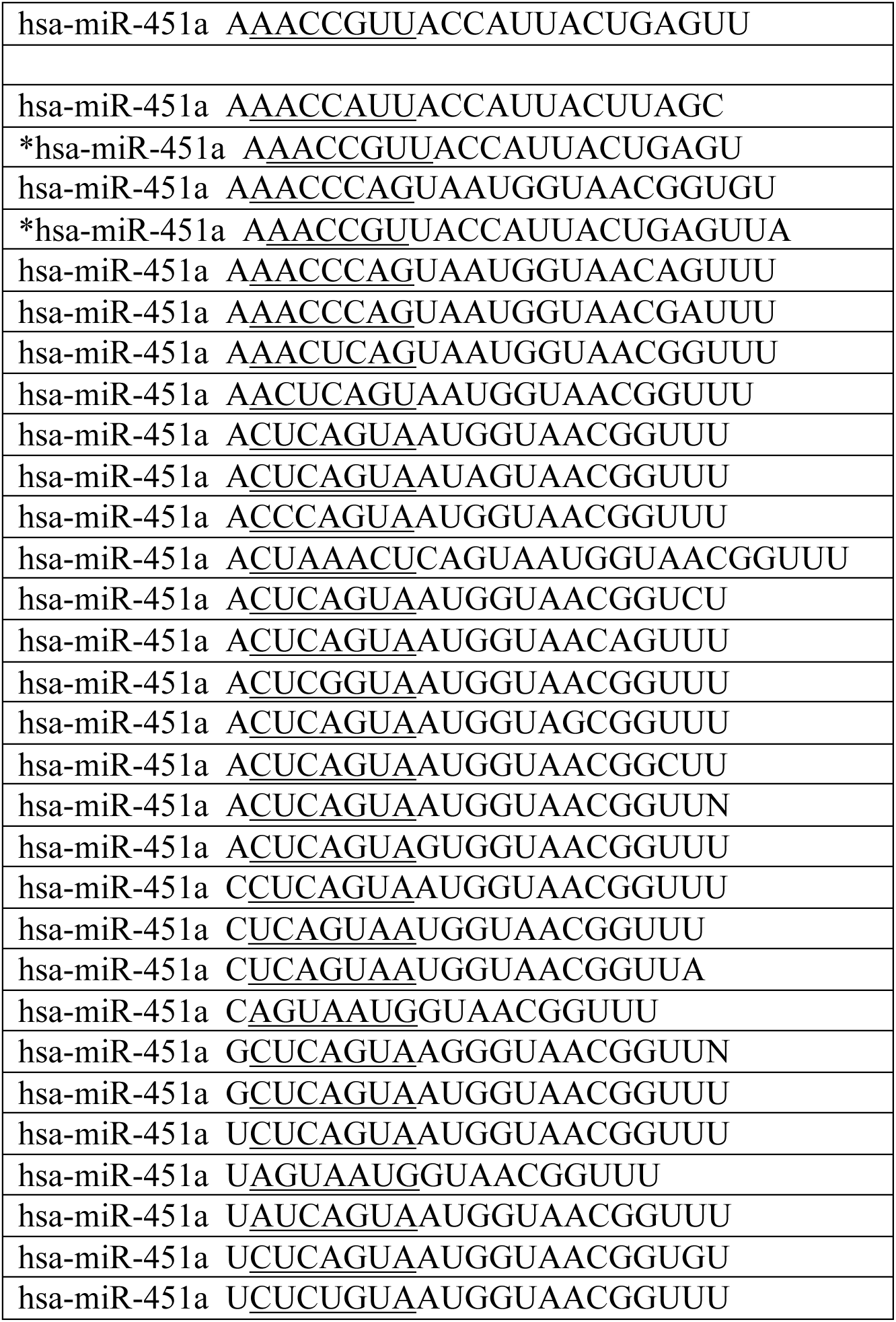
RBC isomiRs. Asterisks labeled isomiRs have the same core sequence as the parent miR451-a.

## Discussion

We here describe a new application of molecular beacons aimed at identification of discreet modification in the length of targets methylation of hydroxy-methylation profile of specific nucleic acid sequence, in addition to insertions and deletions, along with a sample preparation approach, which allows identification of target RNA fragments in a mix and read type approach.

Cell-free nucleic acids are at the core of liquid biopsy, a new method for obtaining disease-relevant samples in small volume^(22)^. Cell-free nucleic acids is a rather an inaccurate, umbrella term encompassing in addition to genomic and mitochondrial DNA, and over 15 species of RNA including mRNA, t-RNA, circular RNA, PIWI, lnc-RNA and miRNAs^(22-23)^. MicroRNAs are short, non-coding RNA sequences (19-22 nt) that primarily function as silencers of RNA expression, and regulators of gene expression^(24)^. Obtaining relevant information about the presence, amount and PTxMs in target RNA species is currently mired by the requirement of rather large amount of starting material, usually in the microgram range, involved isolation and purification methods followed by extended bioinformatic analyses. We here introduced a sample preparation method comprising of a molecular crowding agent (**Fig. 1)** together with a sample lysis buffer (**Fig. 5**), which act together by releasing the nucleic acid molecules in a conformation suitable for hybridization and increasing the efficacy of the binding several folds. This readout is fully applicable to identifying various ssRNA and ssDNA molecules found in cells or associated with extracellular vesicles. We have previously shown that bead attached MBs capture their intended targets in solution, which then can be analyzed by either flow cytometry or gel electrophoresis^(8)^. This approach may potentially increase the sensitivity of the method even for targets with low copy number, which would otherwise require loading on one lane unrealistic volumes of sample material. The current methylation-sensitive design of the MB does not allow differentiation between the wt and the methylated nucleobases based on fluorescence quantification readouts such as fluorometry or flow cytometry (data not shown), as the fluorochrome exhibits the same fluorescence spectrum regardless of the presence of additional modifications on the nucleobase or ribose/deoxyribose backbone. Gel electrophoresis bypasses this limitation by relying on the subtle differences in the 3-D conformation of the hybrids promoted by the PTxM on the electrophoretic pattern.

The electrophoretic mobility of the MB-miRNAs complex is also sensitive to slight variation in the length of the target, as we have shown in **Fig. 5**. IsomiRs differ from the parent miRNA by several more or fewer nucleotides to either the 3’ (3’ isomiRs), 5’ (5’ isomiRs) end, mixed 3’ and 5’, or by undergoing A-I editing events mediated by ADAR proteins^(17)^. In addition to changing the affinity of the m6A-miRNAs to the target, the newly formed m6A nucleobase has destabilizing effects on the RNA duplex (m6A-switch), preventing the binding of ADAR enzymes to RNA, and consequently the A to I editing rate^(25)^. While the 5’ changes in length are common in 5-15% of the miRNAs, the 3” end edits may happen, depending on the cell type and tissue of origin, to up 50% of a given miRNA specie. Although less common, the changes at the 5’ end of the miRNAs, where the heptamer seed sequence is located, extending from the second to the eight nucleotide, could be more functionally relevant, by targeting new transcripts, different those canonically attributed to the parent miRNA (targetome shifting). If the resulting isomiRs have several added nucleotides outside the binding area, the MB will still bind the target, resulting in hybrid which will migrate slower than the parent miRNA making it difficult to differentiate between them and the parent miRNA with hydroxy-C modification on nucleobases. Conversely, deletion or editing of several nucleotides from either end of the target, will prevent MB binding, rendering a negative signal(^8, 26^). As the databases focused on isomiRs become widespread, and more populated with isomiR species both, in normal and pathological conditions (Tumor IsomiR Encyclopedia, https://isomir.ccr.cancer.gov/browse), designing MBs tailored to the disease relevant isomiRs in addition to the parent miRNAs, may allow differentiation between the two possibilities. Our results presented in **Fig. 5** show that MB synthetized against the parent miR-451 identified 3 isomiRs with different number of nucleotides in RBCs and RBC-derived EVs. The rest of the isomiR-451 detected by RNA-seq in RBC (see Table 3) were not identified by the standard MBs-451a, likely due to their high variability in the sequence, and the single nucleotide specificity of the MB detection, as we and other have shown(^8, 13, 20, 27^). MicroRNA-451a is an RBC-enriched miRNA, which was reported to be involved in protecting the host against *P. falciparum* infection, especially in sickle cell patients, where their miR-451 along with let-7i RBC expression levels are significantly increased. Once the merozoites released by the hepatocytes invade the circulating RBCs the two microRNAs translocate from the RBC cytosol into the parasitophorus vacuole, acting directly on the parasite mRNAs impairing its growth^(28)^. It is conceivable that individuals with mutations in the seed sequence or with low expression levels of certain isomiR-451 or let-7i could be less effective in hindering malaria infection. While miR-451a is primarily described as an RBC-enriched miRNA, its increased expression levels in tumor cells and tumor-derived EVs were shown to limit the invasiveness and cell division rates, in a wide range of cancers ranging from melanoma, lung, esophageal to colon, ovarian and head and neck (reviewed in ^(29)^). In the case of melanoma, the absence of the beneficial effect of miR451 was attributed to the loss of isomiR 451a.1 and decrease expression of the parent miR451.

In our simulation, we observed that the molecular beacon with FL2/Q2 exhibited multiple stable conformations, whether or not the target was present, while FL1/Q1 only resulted in a single stable conformation. This can be attributed to the significantly larger size of FL2/Q2 compared to FL1/Q1. Additionally, we calculated the number of rotatable bonds in each fluorochrome and quencher, with FL2/Q2, and FL1/Q2, having 21, 17, 12, and 13 rotatable bonds, respectively. A larger size and higher number of rotatable bonds can confer greater flexibility to the structure, leading to a more rugged free energy landscape. This, in turn, increases the likelihood of local minima being separated by energy barriers. Each local minimum corresponds to a stable or meta-stable conformation, which explains why FL2/Q2 are more likely to exhibit multiple stable conformations. We are currently utilizing the General AMBER Force Field (GAFF) ^(30)^ for both the fluorochrome and the quencher.

However, the delocalized π-electron orbitals present in numerous fluorescent dyes result in high electronic polarizability, which poses a significant challenge to point charge-based force fields. To address this issue, Grubmüller and colleagues^(31)^ developed AMBER-DYES, a modular fluorescent label force field designed for 22 fluorescent dyes and their corresponding linkers encompassing the Alexa, Atto, and Cy families, which are suitable for several superresolution microscopy techniques. Nevertheless, the fluorescent dyes utilized in our research are not included in the AMBER-DYES force field, therefore, our next step is to create custom parameters for FL1 and FL2 to further improve accuracy.

An ideal PTxM-targeting MB would only fluoresce in the presence of modified nucleobases, allowing a ratiometric determination of the modification percentage of a given nucleic acid in a sample. This, together with the “*mix and read*” lysis buffer would allow multiplex analyses of targets from a wide range of using flow cytometry in formats amenable to clinical settings. Using ML-aided design of the beacons or similar nucleic acid probes, will bypass the need of blind, wet lab testing for the ideal combinations of modified nucleobase and/or backbone with the most suitable fluorochrome and quencher pair. While the method presented here is not suitable to discoveries on new methylation sites on unknown samples, it may however provide a rapid readout of identifying the presence of PTxMs on already known targets, while using reagents and instrumentation available in most research labs. With the help of the ML-assisted testing, designing optimal MB-fluorochrome combinations can be tuned to a wide set of condition-specific targets and unique PTxMs.

## References

1. Zou Y, Mason MG, Wang Y, Wee E, Turni C, Blackall PJ, Trau M, Botella JR. Nucleic acid purification from plants, animals and microbes in under 30 seconds. PLoS Biol. 2017;15(11):e2003916.29161268;PMC5697807

2. Wee SK, Sivalingam SP, Yap EPH. Rapid Direct Nucleic Acid Amplification Test without RNA Extraction for SARS-CoV-2 Using a Portable PCR Thermocycler. Genes (Basel). 2020;11(6);32570810;PMC7349311

3. Cheray M, Etcheverry A, Jacques C, Pacaud R, Bougras-Cartron G, Aubry M, Denoual F, Peterlongo P, Nadaradjane A, Briand J, Akcha F, Heymann D, Vallette FM, Mosser J, Ory B, Cartron PF. Cytosine methylation of mature microRNAs inhibits their functions and is associated with poor prognosis in glioblastoma multiforme. Mol Cancer. 2020;19(1):36;32098627;PMC7041276

4. Li X, Xiong X, Yi C. Epitranscriptome sequencing technologies: decoding RNA modifications. Nature methods. 2016;14(1):23–31;28032622;

5. Helm M, Motorin Y. Detecting RNA modifications in the epitranscriptome: predict and validate. Nat Rev Genet. 2017;18(5):275–91;28216634;

6. Telonis AG, Loher P, Jing Y, Londin E, Rigoutsos I. Beyond the one-locus-one-miRNA paradigm: microRNA isoforms enable deeper insights into breast cancer heterogeneity. Nucleic Acids Res. 2015;43(19):9158–75;26400174;PMC4627084

7. Matsumoto S, Sugimoto N. New Insights into the Functions of Nucleic Acids Controlled by Cellular Microenvironments. Top Curr Chem (Cham). 2021;379(3):17;33782792;

8. Oliveira-Jr GP, Barbosa RH, Thompson L, Pinckney B, Murphy-Thornley M, Lu S, Jones J, Hansen CH, Tigges J, Wong WP, Ghiran IC. Electrophoretic mobility shift as a molecular beacon-based readout for miRNA detection. Biosens Bioelectron. 2021;189:113307;34062334;PMC8461749

9. Liu P, Tian W. Identification of DNA methylation patterns and biomarkers for clear-cell renal cell carcinoma by multi-omics data analysis. PeerJ. 2020;8:e9654;32832275;PMC7409785

10. Jiang X, Liu B, Nie Z, Duan L, Xiong Q, Jin Z, Yang C, Chen Y. The role of m6A modification in the biological functions and diseases. Signal Transduct Target Ther. 2021;6(1):74;33611339;PMC7897327

11. Fang Z, Mei W, Qu C, Lu J, Shang L, Cao F, Li F. Role of m6A writers, erasers and readers in cancer. Exp Hematol Oncol. 2022;11(1):45;35945641;PMC9361621

12. Luitz MP, Barth A, Crevenna AH, Bomblies R, Lamb DC, Zacharias M. Covalent dye attachment influences the dynamics and conformational properties of flexible peptides. PLoS One. 2017;12(5):e0177139;28542243;PMC5441599

13. Tyagi S, Kramer FR. Molecular beacons: probes that fluoresce upon hybridization. Nat Biotechnol. 1996;14(3):303–8;9630890;

14. Kuo WP, Tigges JC, Toxavidis V, Ghiran I. Red Blood Cells: A Source of Extracellular Vesicles. Methods in molecular biology. 2017;1660:15–22;28828644;

15. Muroya T, Kannan L, Ghiran IC, Shevkoplyas SS, Paz Z, Tsokos M, Dalle Lucca JJ, Shapiro NI, Tsokos GC. C4d deposits on the surface of RBCs in trauma patients and interferes with their function. Crit Care Med. 2014;42(5):e364–72;24448198;4099476

16. Worpenberg L, Paolantoni C, Roignant JY. Functional interplay within the epitranscriptome: Reality or fiction? Bioessays. 2022;44(2):e2100174;34873719;

17. Tomasello L, Distefano R, Nigita G, Croce CM. The MicroRNA Family Gets Wider: The IsomiRs Classification and Role. Front Cell Dev Biol. 2021;9:668648;34178993;PMC8220208

18. Loher P, Londin ER, Rigoutsos I. IsomiR expression profiles in human lymphoblastoid cell lines exhibit population and gender dependencies. Oncotarget. 2014;5(18):8790–802;25229428;PMC4226722 interests.

19. Karlsen TA, Aae TF, Brinchmann JE. Robust profiling of microRNAs and isomiRs in human plasma exosomes across 46 individuals. Scientific reports. 2019;9(1):19999;31882820;PMC6934752

20. Manganelli R, Tyagi S, Smith I. Real Time PCR Using Molecular Beacons : A New Tool to Identify Point Mutations and to Analyze Gene Expression in Mycobacterium tuberculosis. Methods in molecular medicine. 2001;54:295–310;21341083;

21. Tyagi S, Bratu DP, Kramer FR. Multicolor molecular beacons for allele discrimination. Nat Biotechnol. 1998;16(1):49–53;9447593;

22. Pos O, Biro O, Szemes T, Nagy B. Circulating cell-free nucleic acids: characteristics and applications. Eur J Hum Genet. 2018;26(7):937–45;29681621;PMC6018748

23. Szilagyi M, Pos O, Marton E, Buglyo G, Soltesz B, Keseru J, Penyige A, Szemes T, Nagy B. Circulating Cell-Free Nucleic Acids: Main Characteristics and Clinical Application. Int J Mol Sci. 2020;21(18);32957662;PMC7555669

24. Bushati N, Cohen SM. microRNA functions. Annu Rev Cell Dev Biol. 2007;23:175–205

25. Xiang JF, Yang Q, Liu CX, Wu M, Chen LL, Yang L. N(6)-Methyladenosines Modulate A-to-I RNA Editing. Mol Cell. 2018;69(1):126–35 e6;29304330;

26. Giesendorf BA, Vet JA, Tyagi S, Mensink EJ, Trijbels FJ, Blom HJ. Molecular beacons: a new approach for semiautomated mutation analysis. Clin Chem. 1998;44(3):482–6;9510851;

27. Marras SAE, Russell Kramer F, Tyagi S. Multiplex detection of single-nucleotide variations using molecular beacons. Genetic Analysis: Biomolecular Engineering. 1999;14(5-6):151–6

28. LaMonte G, Philip N, Reardon J, Lacsina JR, Majoros W, Chapman L, Thornburg CD, Telen MJ, Ohler U, Nicchitta CV, Haystead T, Chi JT. Translocation of sickle cell erythrocyte microRNAs into Plasmodium falciparum inhibits parasite translation and contributes to malaria resistance. Cell host & microbe. 2012;12(2):187–99;22901539;PMC3442262

29. Bai H, Wu S. miR-451: A Novel Biomarker and Potential Therapeutic Target for Cancer. Onco Targets Ther. 2019;12:11069–82;31908476;PMC6924581

30. Sprenger KG, Jaeger VW, Pfaendtner J. The general AMBER force field (GAFF) can accurately predict thermodynamic and transport properties of many ionic liquids. J Phys Chem B. 2015;119(18):5882–95;25853313;

31. Graen T, Hoefling M, Grubmuller H. AMBER-DYES: Characterization of Charge Fluctuations and Force Field Parameterization of Fluorescent Dyes for Molecular Dynamics Simulations. J Chem Theory Comput. 2014;10(12):5505–12;26583233;

